# Effects of graphene oxide and graphite on soil bacterial and fungal diversity

**DOI:** 10.1101/530485

**Authors:** Christian Forstner, Thomas G. Orton, Adam Skarshewski, Peng Wang, Peter M. Kopittke, Paul G. Dennis

## Abstract

Graphene oxide (GO) is an oxidized form of graphene that is relatively cheap and easy to produce. This has heralded its widespread use in a range of industries, with its likelihood of release into the environment increasing accordingly. In pure culture, GO has been shown to influence bacteria and fungi, but its effects on environmental microbial communities remain poorly characterized, despite the important ecosystem services that these organisms underpin. Here, we characterized the effects of GO and graphite, over time and at three concentrations (1 ng, 1 μg and 1 mg kg dry soil^−1^), on soil bacterial and fungal diversity using 16S rRNA and ITS2 gene amplicon sequencing. Graphite was included as a reference material as it is widely distributed in the environment. Neither GO or graphite had significant effects on the alpha diversity of microbial communities. The composition of bacterial and fungal communities, however, was significantly influenced by both materials at all doses. Nonetheless, the effects of GO and graphite were of similar magnitude, albeit with some differences in the taxa affected.

## 1. Introduction

Graphene oxide (GO) is a nanomaterial with unique properties that has a wide-range of uses (Novoselov et al., 2012; Yoo et al., 2014). The environmental release of GO is predicted to increase along with the concomitant need to understand its potential ecological effects (Ahmed and Rodrigues, 2013). Studies based on pure bacterial and fungal cultures indicate that GO can elicit both positive and negative effects on cell integrity and function. For example, GO can kill or suppress bacterial (Akhavan and Ghaderi, 2010; Liu et al., 2011) and fungal cultures (Chen et al., 2014; Li et al., 2015), by inducing oxidative stress (Ahmed and Rodrigues, 2013; Liu et al., 2011) or penetrating cell-walls and extracting phospholipids (Tu et al., 2013). On the other hand, some studies have shown that GO can act as a respiratory electron acceptor (Salas et al., 2010; Wang et al., 2011) and promote the growth of bacterial cultures (Ruiz et al., 2011). These studies highlight that GO has the potential to affect soil microbial communities, which play fundamental roles in the provision of ecosystem goods and services (Bardgett and van der Putten, 2014; Wagg et al., 2014). Despite this, the impacts of GO on soil microbial diversity and function are poorly understood.

To the best of our knowledge, there are only four previous studies that have directly investigated the effects of GO on soil microbial communities (Chung et al., 2015; Du et al., 2015; Kim et al., 2018; Xiong et al., 2018). The first, focused exclusively on soil microbial biomass and enzyme activities (Chung et al., 2015). It revealed significant, but temporary, effects on the activities of xylosidase, β-1,4-N-acetyl glucosaminidase, and acid phosphatase, and no effects on microbial biomass (Chung et al., 2015). The second study aimed to compare the diversity of bacterial communities in control and GO-amended soils using phylogenetic marker gene sequencing (Du et al., 2015). The authors claimed that GO increased soil bacterial diversity and altered community composition (Du et al., 2015). Nonetheless, as highlighted by Forstner et al. (2016), their claims were unjustified as their study lacked replication within treatments (n=1), and did not account for differences in the numbers of sequences per sample when comparing the numbers of taxa present. The third study investigated the effects of GO on microbial enzyme activities and bacterial diversity in cadmium contaminated soil (Xiong et al., 2018). It revealed that the activities of catalase and dehydrogenase increased, while that of urease decreased in response to GO (Xiong et al., 2018). Furthermore, the composition of bacterial communities, but not their diversity, appeared to be influenced by GO, although no statistical support for this was provided (Xiong et al., 2018). Lastly, the fourth study focused on the effects of silver-GO nanocomposites on soil bacterial communities, but included a GO treatment (Kim et al., 2018). Unfortunately, this GO treatment was not compared to a water-only control; therefore, it was not possible to isolate GO effects from those attributable to Ag (Kim et al., 2018).

Importantly, all previous studies applied GO at doses that are considerably higher than those likely to be encountered in the environment. The main routes of entry for many nanomaterials into the environment are thought to be wastewater treatment and landfill (Gottschalk et al., 2009; Mueller and Nowack, 2008; Sun et al., 2014). While data are not available for GO, models estimate that 3.7-7.1 ng and 0.8-1.6 μg of carbon nanotubes (CNTs) kg soil^−1^ enter European natural and sludge-amended soils every year (Sun et al., 2014). Model estimates for the rate of release of other nanomaterials into soils (nanosilver, nano-ZnO and fullerenes) are similar (Sun et al., 2014). Chung et al. (2015) applied 0.1-1.0 g GO kg soil^−1^, while Xiong et al. (2018) and Du et al. (2015) applied 1.0-2.0 g and 5 g GO kg soil^−1^, respectively. These doses are equivalent to between 62.5 thousand and 1.25 billion times the estimated annual rate of accumulation for CNTs (Sun et al., 2014). Hence, the effects of GO on soil microbial diversity are poorly understood, particularly at concentrations approximating those deemed realistic for similar nanomaterials. In addition, as previous studies have only considered bacterial communities, the effects of GO on other important groups of soil organisms, such as fungi, are not known.

Finally, when characterizing the effects of nanomaterials on soil microbial diversity it is important to include reference materials that provide a context with which to interpret changes (Petersen, 2015). For example, while a significant shift in microbial diversity may be observed in response to a nanomaterial, the magnitude of such a change may be small in comparison to those associated with material that are already widely distributed in the environment. Clearly, therefore, this information is essential for the development of appropriate policy frameworks for the safe and sustainable use of nanomaterials. Despite this, such reference materials were not included in any of the previous studies concerning the effects of GO on soil microbial communities (Chung et al., 2015; Du et al., 2015; Kim et al., 2018; Xiong et al., 2018).

In this study, we investigated the effects of one-off applications of GO and graphite on soil bacterial and fungal diversity using high-throughput phylogenetic marker gene sequencing. Graphite was included as a reference compound as it chemically similar to GO, without being classed as a nanomaterial, and is widely distributed in the environment. GO and graphite were applied at three concentrations (1 ng, 1 μg and 1 mg GO kg soil^−1^) representing environmentally relevant low, high and extremely high rates of release based on those predicted for CNTs (Sun et al., 2014). All treatments were replicated three times and communities were characterized 7, 14 and 30 days after application.

## 2. Materials and methods

### 2.1 Experimental design

The soil used in this study is classified as a Kandosol according to the Australian Soil Classification (Isbell, 2002), or an Ultisol according to the USDA Soil Taxonomy (Soil Survey Staff, 2014) and has been described in our previous work (see Table S1 from Wang *et al*. (2016)). Briefly, the soil was collected at a depth of 0–20 cm from a pineapple (*Ananas comosus*) farm in Queensland, Australia. The soil had a sandy loam texture, pH of 5.4 (1:5 soil/water), conductivity of 0.1 dS/m (1:5 soil/water), and total organic Carbon content of 1.1%. Approximately 11 kg of fresh soil was passed through a 2 mm sieve and then split into seven c. 1.6 kg sub-samples to which the treatments were applied. Graphene oxide (Aldrich Chemistry, CAT: 763705, Lot# MKBQ8029V) and graphite (Sigma Aldrich, CAT:282863, size <20 μm) were applied to soils at rates of 1 ng, 1 μg and 1 mg kg dry soil^−1^ using a fine mist sprayer and thoroughly homogenized by mechanical mixing. In brief, either the GO or graphite was dispersed into deionized water at a volume that resulted in the soil being adjusted to 50% water holding capacity (WHC). The solution was then sprayed on the soil prior to mechanical mixing. These three rates were selected to represent environmentally relevant, high and extremely high concentrations based on release rates predicted for CNTs (Sun et al., 2014). Soils in the control treatments were sprayed with an equal quantity of deionized water only and then mixed in the same manner as for the GO and graphite treatments. Three replicate 500 g samples of each treatment were placed into 1 L plastic containers with lids that facilitated gas exchange. This yielded 21 containers that were incubated for 30 days in the dark at 25°C, with the humidity maintained at 80% in order to keep the soils at the same WHC throughout the experimental period.

### 2.2 Soil sampling and DNA extraction

Soil cores, of approximately 25 g, were collected from each experimental unit after 7, 14 and 30 days using sterile 50 ml plastic tubes, and immediately transferred to −80°C storage. DNA was extracted from 250 mg of thawed soil using the Power Soil DNA Isolation kit (MO BIO Laboratories, Carlsbad, CA) according to the manufacturers’ instructions. To avoid systematic biases we randomized the order of samples for all processing steps.

### 2.3 PCR amplification and sequencing of phylogenetic marker genes

Universal bacterial 16S rRNA genes were amplified by polymerase chain reaction (PCR) using the primers 926F (5’-AAA CTY AAA KGA ATT GRC GG-3’) (Engelbrektson et al., 2010) and 1392wR (5’-ACG GGC GGT GWG TRC-3’) (Engelbrektson et al., 2010). Fungal ITS2 regions were PCR amplified using the primers gITS7F (5’- GTG ART CAT CGA RTC TTT G -3’) (Ihrmark et al., 2012) and ITS4R (5’- TCC TCC GCT TAT TGA TAT GC -3’) (White et al., 1990). For both groups, the forward and reverse primers were modified on the 5’ end to contain the Illumina overhang adapter for compatibility with the i5 and i7 Nextera XT indices, respectively. PCRs were performed with 1.5 μl template DNA, in 1X PCR Buffer minus Mg^2+^ (Invitrogen), 100 μM of each of the dNTPs (Invitrogen), 300 μM of MgCl2 (Invitrogen), 0.625 U Taq DNA Polymerase (Invitrogen), and 250 μM of each primer, made up to a total volume of 25 μl with molecular biology grade water. Thermocycling conditions were as follows: 94°C for 3 minutes; then 35 cycles of 94°C for 45 seconds, 55°C for 30 seconds, 72°C for 1 minute 30 seconds; followed by 72°C for 10 minutes. Amplifications were performed using a Veriti^®^ 96-well Thermocycler (Applied Biosystems). PCR success was determined by gel electrophoresis, which also facilitated visual confirmation of amplicon size and quality.

Amplicons were purified using AMPure magnetic beads (Agencourt) and subjected to dual indexing using the Nextera XT Index Kit (Illumina) as per the manufacturer’s instructions. Indexed amplicons were purified using AMPure XP beads and then quantified using a PicoGreen dsDNA Quantification Kit (Invitrogen). Equal concentrations of each sample were pooled and sequenced on an Illumina MiSeq at The University of Queensland’s Institute for Molecular Biosciences (UQ, IMB) using 30% PhiX Control v3 (Illumina) and a MiSeq Reagent Kit v3 (600 cycle; Illumina) according the manufacturer’s instructions.

### 2.4 Processing of sequence data

Data were analyzed using a modified UPARSE pipeline (Edgar, 2013). For both datasets, analyses were performed using the forward reads only. For 16S rDNA sequences, USEARCH (v10.0.240) (Edgar, 2010) was used to perform the following steps: 1) primers were removed and the residual sequences were trimmed to 250 bp using fastx_truncate; 2) high-quality sequences were identified using fastq_filter by discarding reads with greater than one expected error (-fastq_maxee=1); 3) duplicate sequences were removed using fastx_uniques; 4) sequences were clustered at 97% similarity into operational taxonomic units (OTU) and potential chimeras were identified and removed using cluster_otus; and 5) an OTU table was generated using otutab with default parameters from the pre-trimmed reads and the OTU representative sequences. For the ITS data, ITSx v1.0.11 (Bengtsson-Palme et al., 2013) was used to identify and extract fungal ITS2 sequences. Chimeric ITS2 sequences were identified and removed using the uchime2_ref command of USEARCH and the UNITE database (v7.2 - 2017.10.10) (Nilsson et al., 2019). ITS2 sequences were then clustered at 97% similarity into operational taxonomic units (OTU) and an OTU table was generated using the otutab command of USEARCH with default parameters. SILVA SSU (v128) (Quast et al., 2013) and UNITE (v7.2-2017.10.10) (Nilsson et al., 2019) taxonomy was assigned to the 16S and ITS sequences, respectively, using BLASTN (v2.3.0+) (Zhang et al., 2000) within the feature classifier of QIIME2 (v2017.9) (Boylen et al., 2018). The 16S OTU table was then filtered to remove OTUs classified as chloroplasts, mitochondria, archaea or eukaryotes using the BIOM (McDonald et al., 2012) tool suite. For 16S, de-novo multiple sequence alignments of the representative OTU sequences were generated using MAFFT (v7.221) (Katoh and Standley, 2013) and masked with the alignment mask command of QIIME2. The masked alignment was used to generate a midpoint-rooted phylogenetic tree using FastTree (v2.1.9) (Price et al., 2010) in QIIME2. OTU tables were rarefied to 4850 and 1150 sequences per sample for 16S and ITS, respectively. The mean numbers of observed (Sobs) and predicted (Chao1) OTUs were calculated using QIIME2 for both bacteria and fungi. For bacteria, we also calculated Faith’s phylogenetic diversity index (Faith’s PD) (Faith, 1992) using QIIME2.

### 2.5 Statistical analyses

For statistical analyses, we defined Treatment as the combination of applied substance (none for the control, GO or graphite) and applied dose (1 ng, 1 μg or 1 mg kg dry soil^−1^). Hence, Treatment was defined as a categorical variable with seven classes. In order to determine whether the GO and graphite treatments significantly affected the alpha diversity metrics, we used a linear mixed-effects model approach (Pinheiro and Bates, 2004). Treatment (as defined above) and Day, as well as their interaction, were treated as fixed effects, and soil containers (samples) were treated as a random effect to account for the repeated measures. F-tests were applied to assess significance (*P*<0.05), and were implemented in R using the lme4 (Bates et al., 2015) and lmerTest (Kuznetsova, 2017) packages.

Differences in the relative abundances of taxa between samples (beta diversity), and the inferred functional profiles were assessed using multivariate generalized linear models using a negative binomial distribution (Warton, 2011). The significance of differences in community composition was determined by comparing the sum-of-likelihood test statistics for the alternative statistical models via a resampling method (Wang et al., 2012) that accounted for the correlation between species and the correlation within the repeated measures taken from the same sample container. These comparisons were implemented in R using the mvabund package (Wang et al., 2012). Taxa whose maximum relative abundance was less than 0.1% for bacteria or 0.4% for fungi were disregarded before statistical analysis. Where an interactive effect of Treatment and Day was significant, post-hoc analyses were undertaken to investigate which Treatments differed on what Days. Where no interactive effect was found, but a main effect of Treatment was, post-hoc analyses focused on which Treatments differed from one another. The Benjamini-Hochberg correction was applied to all post-hoc tests.

### 2.6 Data availability

All sequences have been deposited to the Sequence Read Archive (SRA) under BioProject accession number *GenBank: PRJNA515098*.

## 3. Results

### 3.1 Alpha diversity of soil microbial communities

The numbers of observed (Sobs) and predicted (Chao1) bacterial and fungal taxa, as well as the phylogenetic diversity of bacterial communities (Faith’s PD), were not significantly influenced by any of the GO or graphite treatments relative to the controls throughout the experiment (Fig. 1, S1).

**Fig. 1.**
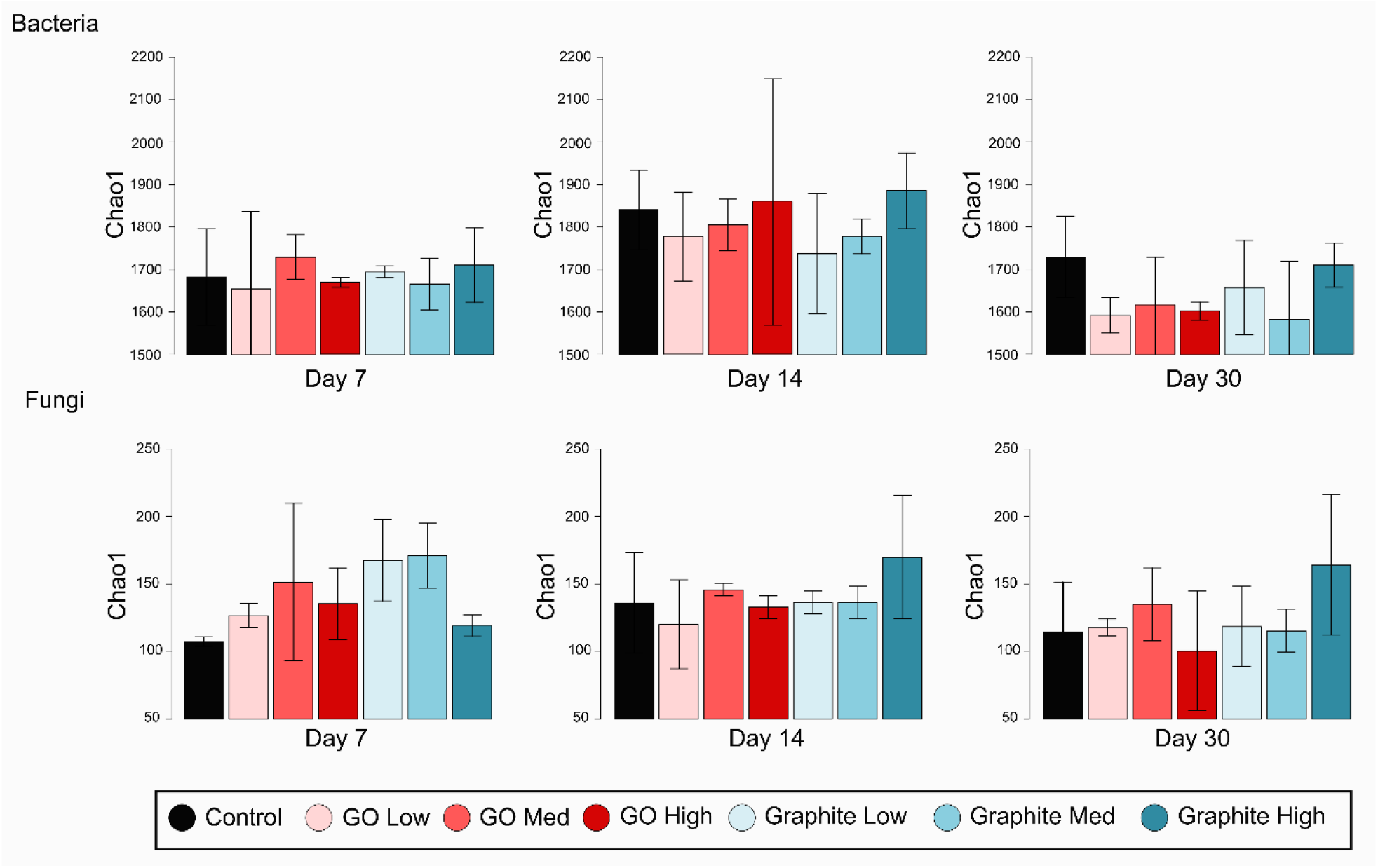
The numbers of predicted bacterial and fungal OTUs (Chao1) after 7, 14 and 30 days by treatment. The error bars represent standard deviations. None of the treatments differed significantly from the controls. GO and graphite doses correspond to 1 ng, 1 μg and 1 mg kg^−1^ soil.

### 3.2 Soil bacterial community composition

Relative to the controls, the composition of soil bacterial communities was significantly influenced by the addition of GO and graphite, and these effects differed over time (*P* < 0.001). Effects were observed for both materials at all doses except for the low GO dose on day 14 (Tables 1 and S1). Albeit significant, there was not a consistent direction of change in bacterial community composition with increasing GO or graphite dose (Fig. 2). Bacterial community composition differed significantly between all GO and graphite treatments (Table S2).

**Fig. 2.**
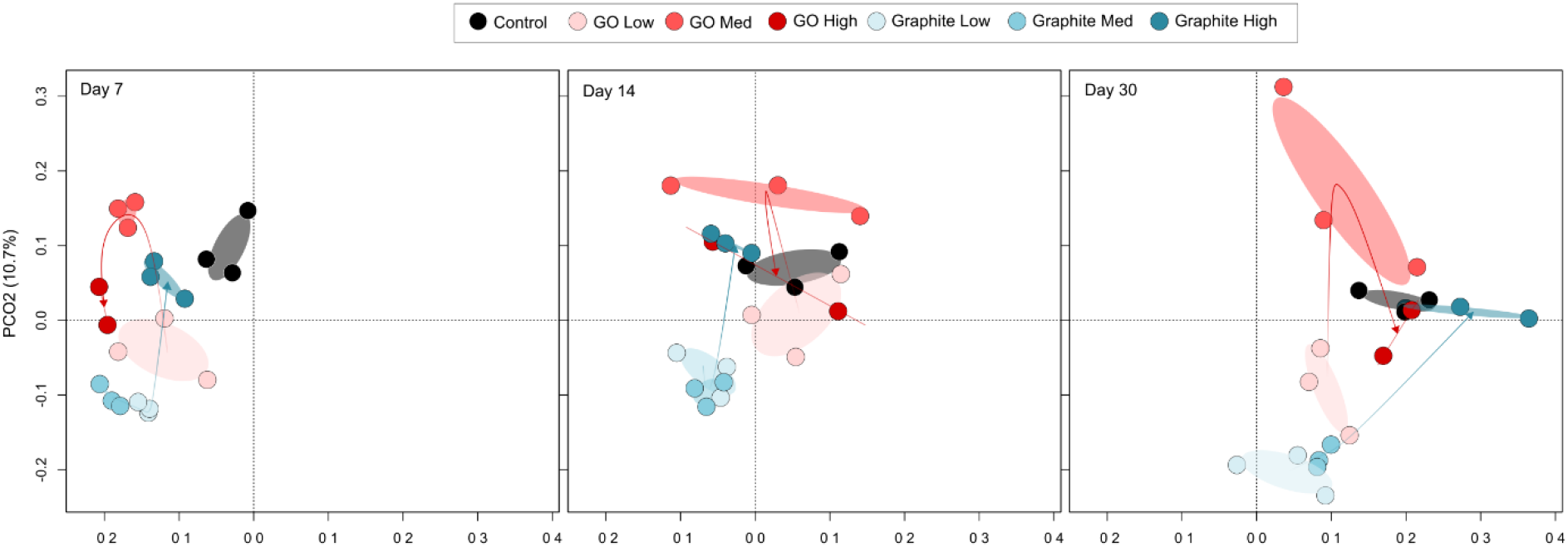
Principal coordinate analysis (PCoA) ordination illustrating differences in the composition of bacterial communities in control, and graphite and GO amended soils over time. The ellipses represent standard deviations. The three panels, representing different time points, derive from a single ordination that included all samples. Each time point is shown separately to avoid over-cluttering. The arrows for each treatment move from low, through medium, to high dose. They highlight that there was not a consistent direction of change in community composition with increasing GO or graphite dose. GO and graphite doses correspond to 1 ng, 1 μg and 1 mg kg^−1^ soil.

**Table 1:** Summary of multivariate GLM post-hoc results computed using mvabund highlighting differences in bacterial community composition between treatments relative to the controls within each time point.

| Test | *P* value |
| --- | --- |
| Day 7 GO Low | <0.001*** |
| Day 7 GO Medium | <0.001*** |
| Day 7 GO High | <0.001*** |
| Day 7 Graphite Low | <0.001*** |
| Day 7 Graphite Medium | <0.001*** |
| Day 7 Graphite High | <0.001*** |
| Day 14 GO Low | 0.140 |
| Day 14 GO Medium | 0.003** |
| Day 14 GO High | 0.020* |
| Day 14 Graphite Low | <0.001*** |
| Day 14 Graphite Medium | <0.001*** |
| Day 14 Graphite High | 0.006** |
| Day 30 GO Low | 0.001** |
| Day 30 GO Medium | <0.001*** |
| Day 30 GO High | 0.021* |
| Day 30 Graphite Low | <0.001*** |
| Day 30 Graphite Medium | <0.001*** |
| Day 30 Graphite High | 0.001** |

Bacterial communities were dominated by representatives of the Acidobacteria, Actinobacteria, Armatimonadetes, Bacteriodetes, Candidatus Berkelbacteria, Chlamydiae, Chlorobi, Chloroflexi, Fibrobacteres, Firmicutes, Gemmatimonadetes, Microgenomates, Planctomycetes and Proteobacteria, Saccharibacteria, and Verrucomicrobia (Fig. S2).

The 100 OTUs that were most strongly associated with differences in community composition between treatments were obtained from the multivariate generalized linear models (GLMs) and assessed independently using univariate GLM models. Of these 49 were found to differ significantly from the control in at least one treatment combination after Benjamini-Hochberg correction for multiple comparisons (Fig. 3). Four OTUs responded exclusively to GO: a member of the *Tepidisphaeraceae* (OTU81, Planctomycetes) and an *Oligoflexales* (OTU363, Deltaproteobacteria) that increased in relative abundance in the presence of GO; a Ktedonobacteria (OTU1084, Chloroflexi) population which declined; and a representative of the *Acidobacteriaceae* (OTU2196, Acidobacteria), which increased at some doses but declined at others (Fig. 3). An additional 30 OTUs, representing a broad-range of phyla, responded to both GO and graphite (Fig. 3). The remaining 15 OTUs, again of broad phylogenetic coverage, responded exclusively to graphite (Fig. 3).

**Fig. 3.**
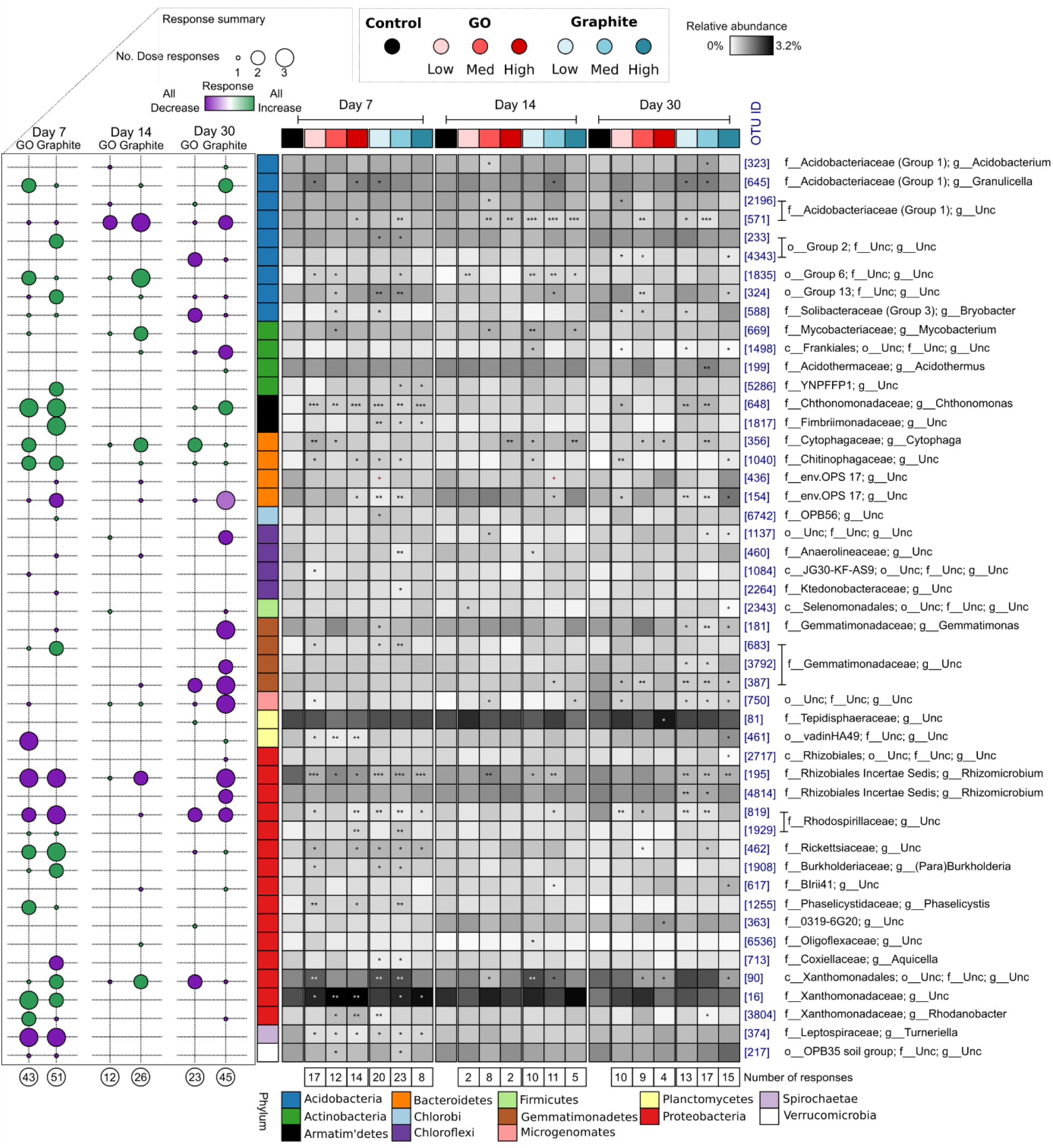
Heatmap of the relative abundances of 49 bacterial OTUs that differed significantly from the control in at least one treatment combination. The asterisks highlight which treatments differ significantly from the controls on each day (*P* < 0.05*, *P* < 0.01**, *P* < 0.001***). Each column of the heatmap represents the mean relative abundance of each treatment (n = 3). The bubble-plot on the left summarizes the number (circle size) of GO or graphite doses that an OTU responded to relative to the controls, and of these how many manifested as increases or decreases in relative abundance (circle color). The numbers below the bubble plot and heatmap show the total numbers of significant responses to a particular treatment relative to the control within the same day. The OTU IDs are consistent throughout the manuscript. The phylum of each OTU is indicated by the colors on the left of the heatmap and the affiliations associated with each color are shown at the bottom. GO and graphite doses correspond to 1 ng, 1 μg and 1 mg kg^−1^ soil.

### 3.3 Soil fungal community composition

Relative to the controls, the composition of soil fungal communities was significantly influenced by the addition of GO and graphite at all doses (*P* = 0.001), and these treatment effects did not differ significantly over time (Tables 2 and S3). As observed for bacteria, there was not a consistent direction of change in the composition of fungal communities with increasing GO or graphite dose (Fig. 4). Fungal community composition differed significantly between all GO and graphite treatments (Table S3).

**Fig. 4.**
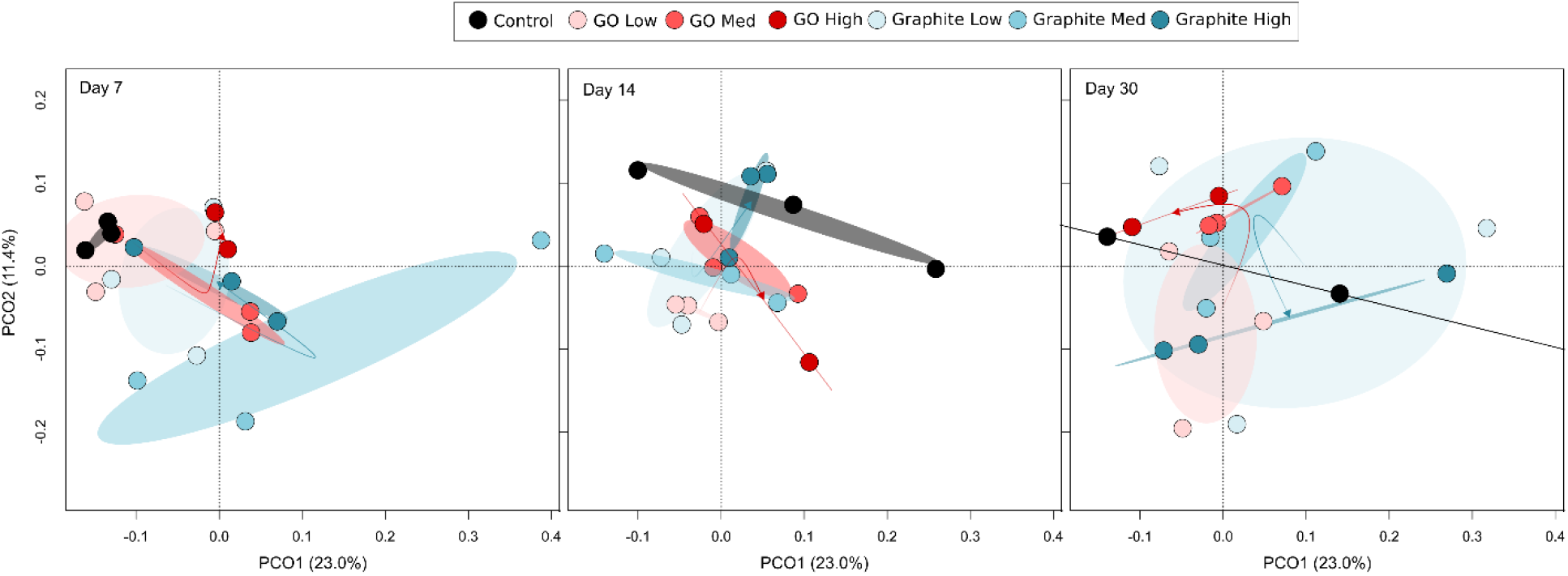
Principal coordinate analysis (PCoA) ordination illustrating differences in the composition of fungal communities in control, and graphite and GO amended soils over time. The ellipses represent standard deviations. The three panels, representing different time points, derive from a single ordination that included all samples. Each time point is shown separately to avoid over-cluttering. On day 30, one of the control samples (PCO1 = 1.2, PCO2 = −0.07) is beyond the plotted area. The arrows for each treatment move from low, through medium, to high dose. They highlight that there was not a consistent direction of change in community composition with increasing GO or graphite dose. GO and graphite doses correspond to 1 ng, 1 μg and 1 mg kg^−1^ soil.

**Table 2:** Summary of multivariate GLM post-hoc results computed using mvabund highlighting differences in fungal community composition between treatments relative to the controls.

| Test | P value |
| --- | --- |
| GO Low | 0.001** |
| GO Medium | 0.005** |
| GO High | 0.005** |
| Graphite Low | 0.002** |
| Graphite Medium | 0.001** |
| Graphite High | 0.001** |

Soil fungal communities were dominated by representatives of the Ascomycota, Basidiomycota, Mortierellomycota, Mucoromycota and Chytridiomycota (Fig. S3). As for the bacteria, the fungal OTUs that were most strongly associated with differences in community composition between treatments were obtained from the multivariate GLMs and assessed independently using univariate GLM models. Of these, 16 OTUs were found to differ significantly from the control in at least one treatment combination after Benjamini-Hochberg correction for multiple comparisons (Fig. 5).

**Fig. 5.**
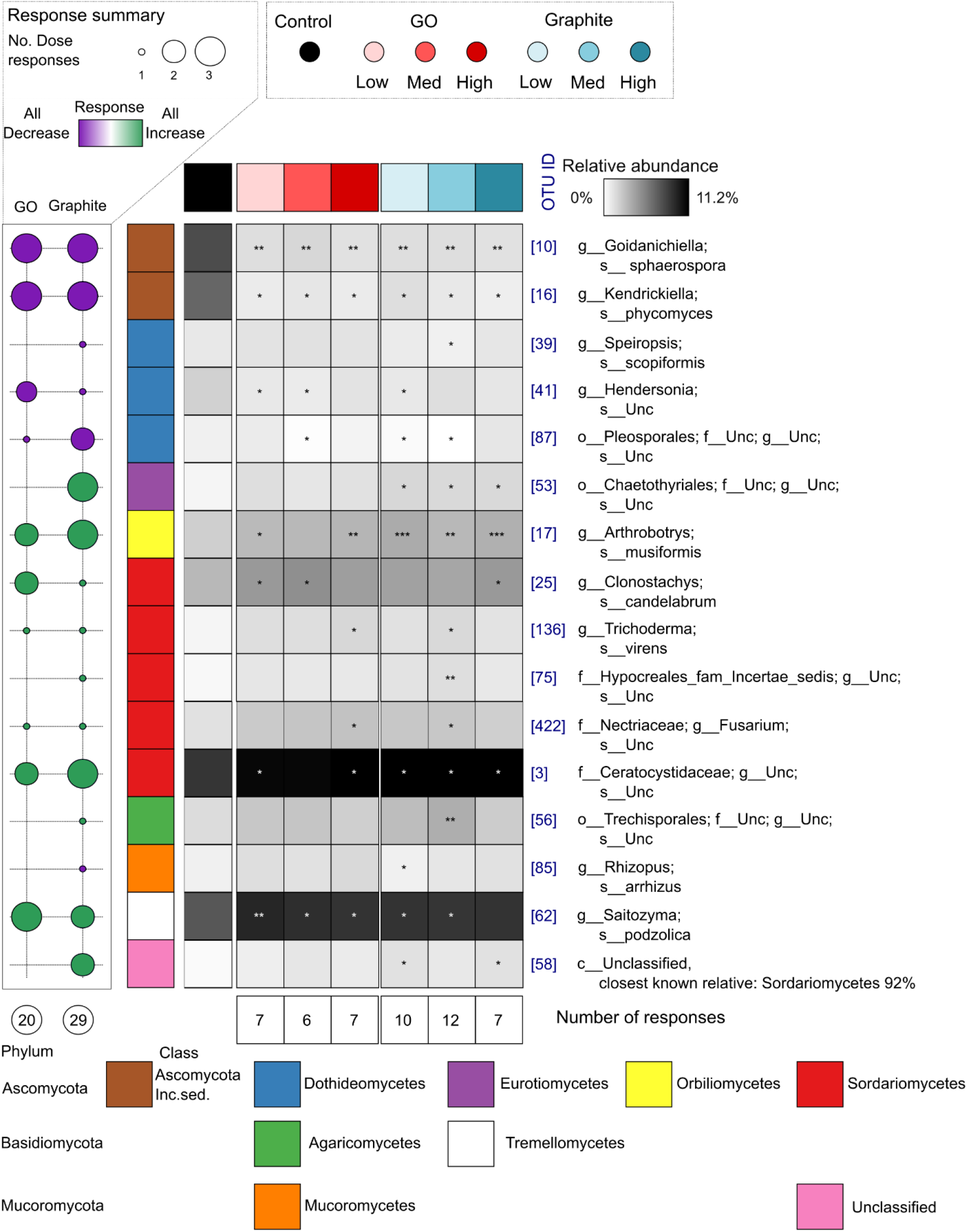
Heatmap of the relative abundances of 16 fungal OTUs that differed significantly from the control in at least one treatment combination. The asterisks highlight which treatments differ significantly from the controls on each day (*P* < 0.05*, *P* < 0.01**, *P* < 0.001***). Each column of the heatmap represents the mean relative abundance of each treatment (n = 9). The bubble-plot on the left summarizes the number (circle size) of GO or graphite doses that an OTU responded to relative to the controls, and of these how many manifested as increases or decreases in relative abundance (circle color). The numbers below the bubble plot and heatmap show the total numbers of significant responses to a particular treatment relative to the control within the same day. The OTU IDs are consistent throughout the manuscript. The phylum and class of each OTU is indicated by the colors on the left of the heatmap and the affiliations associated with each color are shown at the bottom. GO and graphite doses correspond to 1 ng, 1 μg and 1 mg kg^−1^ soil.

No fungal OTUs responded exclusively to GO; however, 10 responded to both GO and graphite (Fig. 5). In all cases the direction of response for these OTUs was the same for GO as it was for graphite. Four of these OTUs were negatively associated with GO and graphite *viz*. a *Goidanichiella sphaerospora* (OTU 10), a *Pleosporales* (OTU 87), a *Kendrickiella phycomyces* (OTU 16) and a *Hendersonia* (OTU 41) population (Fig. 5). In contrast, the relative abundances of six OTUs were positively associated with GO and graphite: an *Arthrobotrys musiformis* (OTU 17), a *Trichoderma virens* (OTU 136), a *Saitozyma podzolica* (OTU 62), a *Ceratocystidaceae* (OTU 3), a *Clonostachys candelabrum* (OTU 25), and a *Fusarium* (OTU 422) population (Fig. 5). The remaining six OTUs responded exclusively to graphite with two decreasing (OTUs 85 and 39) and four increasing (OTUs 53, 56, 75, and 58) in relative abundance, respectively (Fig. 5).

## 4. Discussion

Our study demonstrates that GO can significantly alter the composition, but not alpha diversity, of soil bacterial and fungal communities at loading rates equal to, and beyond, those estimated for the annual accumulation of other nanomaterials (e.g. carbon nanotubes, nanosilver, nano-TiO_2_ and nano-ZnO) in soils (Sun et al., 2014). Nonetheless, our reference material, graphite, also led to significant shifts in community composition, and these were of similar magnitude to those observed for GO, albeit with some differences in the taxa affected.

Of the four previous studies that investigated the effects of GO on soil microbial communities (Chung et al., 2015; Du et al., 2015; Kim et al., 2018; Xiong et al., 2018), three considered bacterial diversity (Du et al., 2015; Kim et al., 2018; Xiong et al., 2018) and two measured microbial enzyme activities (Chung et al., 2015; Xiong et al., 2018). In all cases, the results for bacterial diversity were inconclusive due to a lack of replication (Du et al., 2015), no treatment controls (Kim et al., 2018) or statistical analyses (Du et al., 2015; Xiong et al., 2018). The activities of several enzymes, however, were significantly, albeit only temporally, influenced by GO addition (Chung et al., 2015; Xiong et al., 2018), suggesting that GO has at least some potential to influence soil ecosystem functioning. Nonetheless, all previous studies have focused on the effects of extremely high GO doses – equivalent to between 62.5 thousand and 1.25 billion times the estimated annual rate of accumulation for CNTs (Sun et al., 2014). It could be argued, therefore, that smaller doses would have lesser effects on microbial communities.

Interestingly, our study highlights that the magnitude of community compositional changes observed did not increase with the amount of material applied, or decrease over time. This indicates that even low GO doses (i.e. parts per trillion to parts per billion) can induce community compositional changes that persist beyond 30 days. We speculate that this lack of ‘linear’ dose-response to GO is related to an agglomeration of GO sheets (Dreyer et al., 2010), covering of sheets with other materials (Hui et al., 2014), or interactions with components of the soil matrix (Chen et al., 2018), such as clays which possess a large surface area, capable of binding and immobilizing GO sheets (T. Lu et al., 2017). These interactions could strongly influence the number of GO sheets that interact with soil microbes and significantly alter the properties that they exhibit as pristine nanoparticles. For example, culture-based studies have demonstrated that the edges of GO sheets can cut and destructively extract phospholipids (X. Lu et al., 2017; Tu et al., 2013) from microbial cell membranes. This destructive property of GO is likely to be undermined by interactions that would reduce the number of ‘sharp’ edges (Akhavan and Ghaderi, 2010; Liu et al., 2011).

While there were significant differences in community composition between GO and graphite amended soils, the majority of OTUs that discriminated between treatments, responded in a similar manner to both materials. For example, 61% and 63% of discriminating bacterial and fungal taxa, respectively, responded to both GO and graphite. In contrast, just 8% of discriminating bacterial taxa, and no fungal taxa, responded exclusively to GO; while 31% and 38% of discriminating bacterial and fungal taxa, respectively, responded exclusively to graphite. In general, there were roughly equal numbers of positive and negative responses for taxa responding to GO and/or graphite. This observation corroborates pure culture studies, which report both positive and negative effects of GO on bacterial and fungal isolates (Ahmed and Rodrigues, 2013; Akhavan and Ghaderi, 2010; Chen et al., 2014; Li et al., 2015; Liu et al., 2011; Ruiz et al., 2011; Salas et al., 2010; Tu et al., 2013; Wang et al., 2011). One of the ways in which GO may benefit microbes is by acting as a terminal electron acceptor (Salas et al., 2010; Wang et al., 2011); however, we did not find evidence that respiratory pathways were affected by GO and did not detect any taxa that are well-known to perform extracellular electron transfer. In summary, fewer populations responded exclusively to GO than to graphite; most responded to both materials; and the numbers of positively and negatively affected taxa were about equal.

Despite lacking replication and/or statistical analyses, previous studies that investigated the effects of GO on soil microbial diversity have drawn attention to apparent differences in the relative abundances of ecologically significant taxa, such as those involved in nitrogen cycling. Xiong et al. (2018), for example, reported an increase in the relative abundance of *Rhizobiales* populations and decreases in the relative abundances of *Rhodospirrillaeceae* and *Nitrospirae*. In our study, just one *Rhizobiales* (OTU 195) population was affected by GO addition, and this effect manifested as a decrease in its relative abundance at 7 days, followed by an increase after 14 days. Among the *Rhodospirrilaeceae*, we observed an increase in the relative abundance of OTU 1929, and a decrease in the relative abundance of OTU 819 in response to GO addition (Fig. 3). We did not detect any significant changes in the relative abundances of members of the *Nitrospirae* in response to GO or graphite. To the best of our knowledge, the effects of GO on soil fungal communities have not been previously examined. In pure culture, GO has been shown to suppress the growth of *Fusarium oxysporum*, a species known to contain multiple plant pathogens (Chen et al., 2014). Despite being present within our inventories, we did not detect significant effects of GO on this species; however, the relative abundance of an unclassified relative within the same genus (OTU 422) was observed to increase in response to both GO and graphite addition (Fig. 5).

## 5. Conclusion

As the use of GO increases, its release into the environment will rise. Our study demonstrates that that GO and graphite can influence soil bacterial and fungal community composition, but that their effects are of similar magnitude, albeit with some differences in the taxa affected. In light of this finding, it is important that future studies examine whether GO-induced changes in microbial diversity are likely to undermine the provision of soil ecosystem goods and services.

## Acknowledgements

The authors gratefully acknowledge financial support from The University of Queensland for an Early Career Researcher Award to PGD. CF gratefully acknowledges funding from the Australian Government’s Department of Education and Training in the form of an Australian Government Research Training Program Scholarship administered by The University of Queensland. PMK is the recipient of an Australian Research Council (ARC) Future Fellowship (ARC FT120100277).

