## Supplementary information for "Effects of graphene oxide and graphite on soil bacterial and fungal diversity"

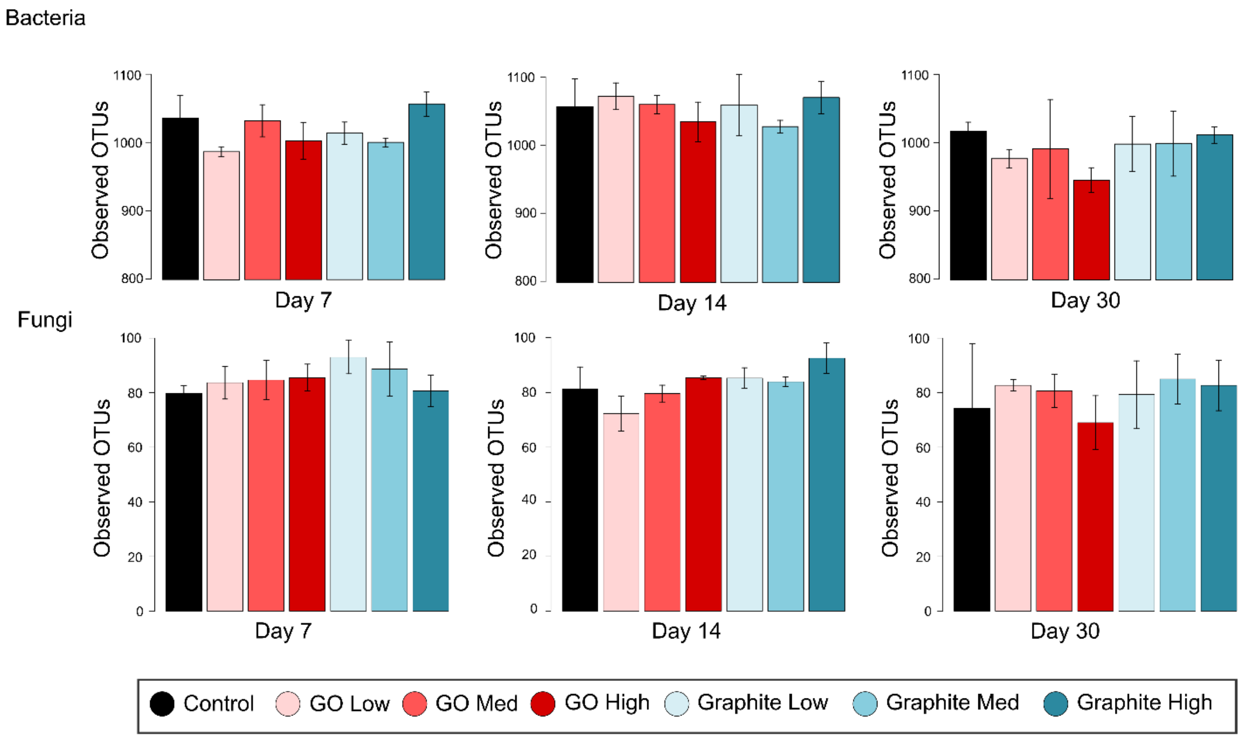

**Fig. S1** The numbers of observed bacterial and fungal OTUs (Sobs) after 7, 14 and 30 days by treatment. The error bars represent standard deviations. None of the treatments differed significantly from the controls. GO and graphite doses correspond to 1 ng, 1 µg and 1 mg $kg^{-1}$ soil.

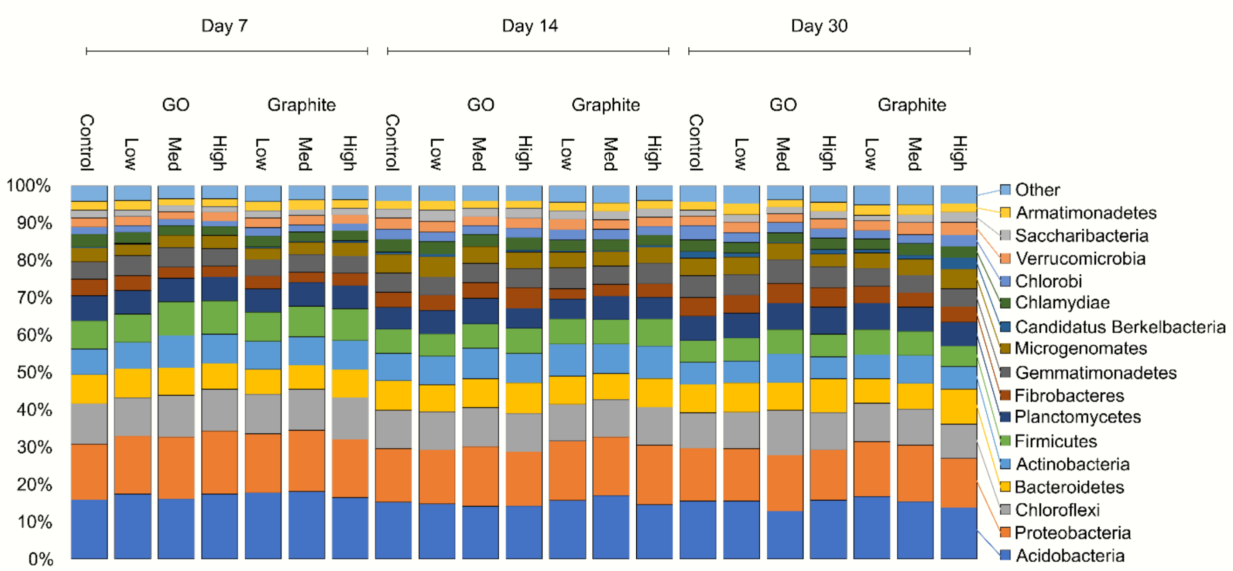

**Fig. S2** The relative abundances of bacteria phyla in control, and GO and graphite amended soils over time. All phyla representing <1% relative abundance are combined as “Other”. GO and graphite doses correspond to 1 ng, 1 μg and 1 mg $kg^{-1}$ soil.

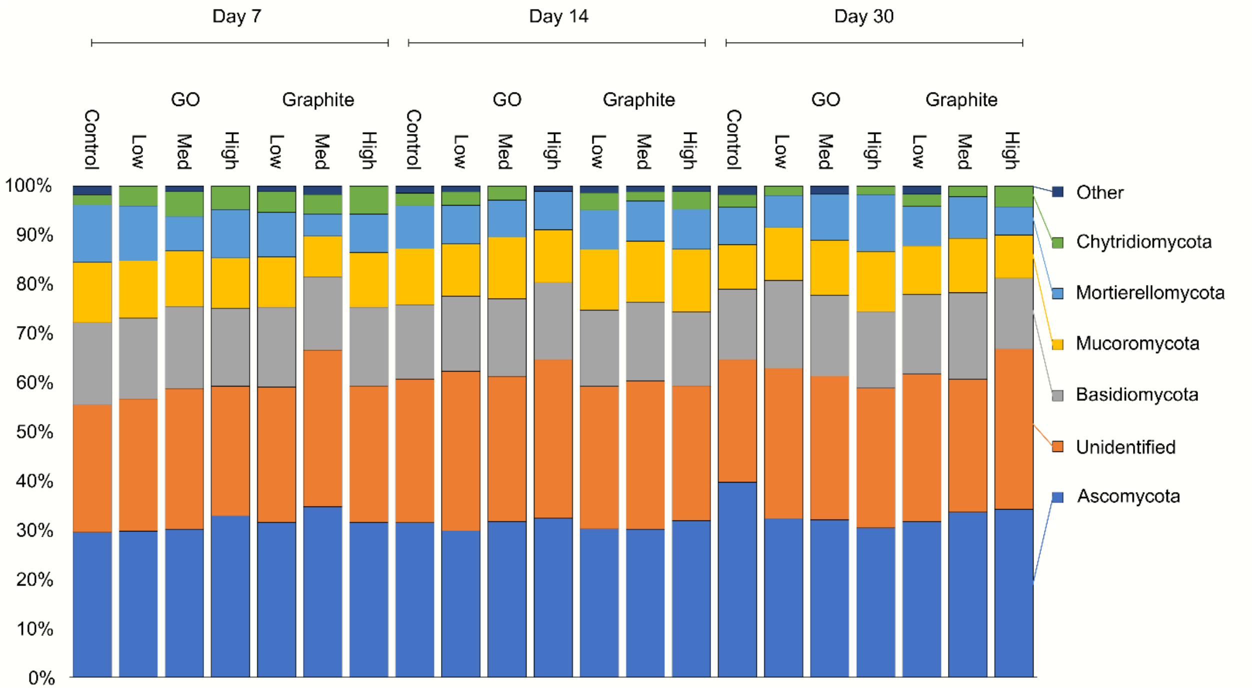

**Fig. S3** The relative abundances of fungal phyla in control, and GO and graphite amended soils over time. All phyla representing <1% relative abundance are combined as “Other”. GO and graphite doses correspond to 1 ng, 1 µg and 1 mg $kg^{-1}$ soil.

**Table S1** Summary of multivariate GLM post-hoc results computed using mvabund highlighting differences in bacterial community composition between difference doses of the same material over time.

| Day | Material | Dose 1 | Dose 2 | *P* value |
|---|---|---|---|---|
| 7 | Graphite | Low | Medium | 0.001 ** |
| 7 | Graphite | Low | High | <0.001 *** |
| 7 | Graphite | Medium | High | <0.001 *** |
| 7 | GO | Low | Medium | <0.001 *** |
| 7 | GO | Low | High | <0.001 *** |
| 7 | GO | Medium | High | <0.001 *** |
| 14 | Graphite | Low | Medium | 0.012 * |
| 14 | Graphite | Low | High | <0.001 *** |
| 14 | Graphite | Medium | High | <0.001 *** |
| 14 | GO | Low | Medium | 0.004 ** |
| 14 | GO | Low | High | 0.702 |
| 14 | GO | Medium | High | 0.006 ** |
| 30 | Graphite | Low | Medium | 0.079 |
| 30 | Graphite | Low | High | <0.001 *** |
| 30 | Graphite | Medium | High | <0.001 *** |
| 30 | GO | Low | Medium | <0.001 *** |
| 30 | GO | Low | High | 0.030 * |
| 30 | GO | Medium | High | 0.009 ** |

**Table S2** Summary of multivariate GLM post-hoc results computed using mvabund highlighting differences in bacterial community composition between GO and graphite at the same dose.

| Day | Dose | Material 1 | Material 2 | *P* value |
|---|---|---|---|---|
| 7 | Low | Graphite | GO | <0.001 *** |
| 7 | Medium | Graphite | GO | <0.001 *** |
| 7 | High | Graphite | GO | 0.007 ** |
| 14 | Low | Graphite | GO | 0.003 ** |
| 14 | Medium | Graphite | GO | <0.001 *** |
| 14 | High | Graphite | GO | 0.035 * |
| 30 | Low | Graphite | GO | 0.009 ** |
| 30 | Medium | Graphite | GO | <0.001 *** |
| 30 | High | Graphite | GO | 0.021 * |

**Table S3** Summary of multivariate GLM post-hoc results computed using mvabund highlighting differences in fungal community composition between difference doses of the same material.

| **Treatment 1** | **Treatment 2** | ***P* value** |
|---|---|---|
| Graphite Low | Graphite Medium | 0.117 |
| Graphite Low | Graphite High | 0.017* |
| Graphite Medium | Graphite High | 0.016* |
| GO Low | GO Medium | 0.005** |
| GO Low | GO High | 0.013* |
| GO Medium | GO High | 0.186 |

**Table S4** Summary of multivariate GLM post-hoc results computed using mvabund highlighting differences in fungal community composition between GO and graphite at the same dose.

| Dose | Material 1 | Material 2 | *P* value |
|---|---|---|---|
| Low | Graphite | GO | 0.011 * |
| Medium | Graphite | GO | 0.011 * |
| High | Graphite | GO | 0.027 * |
